# MelAnalyze: Fact-Checking Melatonin claims using Large Language Models and Natural Language Inference

**DOI:** 10.1101/2024.03.21.586201

**Authors:** Nikitha Karkera, Samik Ghosh, Germaine Escames, Sucheendra K. Palaniappan

## Abstract

With the explosion of health related information in mainstream discourse, distinguishing accurate health-related claims from misinformation is important. Using computational tools and algorithms to help is key. Our focus in this paper is on the hormone Melatonin which is claimed to have broad health benefits and largely sold as a supplement. This paper introduces ‘MelAnalyze,’ a framework for using generative and transformer-based deep learning models adapted as a natural language inference (NLI) task, to semi-automate the fact-checking of general melatonin claims. MelAnalyze is built upon a comprehensive collection of melatonin-related scientific abstracts from PubMed for validation. The framework incorporates components for precise extraction of information from scientific literature, semantic similarity and NLI. At its core, MelAnalyze leverages pre trained NLI models that are fine-tuned on melatonin-specific claims along with semantic search based on vectorized representation of the articles. The best models, fine-tuned on LLaMA1 and RoBERTa, attain good precision, recall, and F1-scores of approximately 0.92. We also introduce a user-friendly web-based tool for fact-checking algorithm evaluation and use. In summary, we show MelAnalyze’s role in empowering users and researchers to assess melatonin-related claims using evidence-based decision-making.

## 1 Introduction

In this era of information overload, it is easier than ever to access and share knowledge. Online platforms and social media networks serve as mediums for the global exchange of such knowledge. While such mediums allow easy dissemination of breakthroughs, health insights etc., they also can be used for spreading of false information. Such false information, especially when related to medical and health areas, can quickly proliferate and even lead to misleading product claims on e-commerce websites for driving sales^1^. These claims often directly influence consumer purchasing behaviors. Traditionally scientific claims were largely limited to academic journals and expert discussions, but nowadays they quickly become a part of public discourse, from traditional remedies’ effectiveness to modern pharmaceuticals’ safety^2^. Their potential impact calls for a systematic approach to validate these claims. This issue is even more a cause of concern when considering natural compounds, such as supplements, which are usually outside the purview regulatory authorities like the FDA. The effects of this misinformation include influencing consumer health choices, misleading consumers on product efficacies, and even potentially leading to adverse health outcomes^3^. Using scientific evidence for validation, we can counter the false claims effectively^4^. However, manually sifting through the vast number of scientific literature to validate every claim is neither feasible nor efficient. Automated fact-checking can be used as a powerful tool to tackle the false information challenge. Recent advancements in Natural Language Processing (NLP) with models such GPT-3, GPT-4 and BERT have led to robust tools for processing and natural language understanding. These models excel in a range of language-related tasks, from machine translation to sentiment analysis, but their potential in fact-checking lies in their capacity to comprehend and contextualize textual information^5,6^.

As a case study to drive our approach for automated fact-checking, we focus on the domain of Melatonin-related claims. Melatonin, is a hormone produced in the pineal gland has seen a lot of interest for its diverse physiological effects, including an important role in circadian rhythms and applications for curing conditions such as sleep disorders and jet lag^**?**^. Melatonin seeems to have a multifaceted role in regulating various physiological processes. Scientific studies have explored its potential as a sleep aid, regulator of circadian rhythms^7–9^, as well antioxidant and anti-inflammatory effects. In fact, public interest in melatonin extends beyond its role in circadian rhythms and sleep regulation. Melatonin has been marketed as a potential treatment for various health conditions, including anxiety, depression, and even cardiovascular diseases, cancer, sepsis and COVID-19^10,11^. A large number of claims surrounding melatonin’s health benefits have surfaced in consumer products. However, alongside legitimate claims, their is considerable misinformation too. Given the strong science based evidence for understanding melatonin, it makes them a good case study for automated fact-checking.

This paper introduces ‘MelAnalyze,’ a computational framework that builds on generative and transformer based deep learning models formalized as a natural language inference (NLI) task to facilitate semi-automatic fact-checking of general melatonin claims. The framework integrates melatonin-related scientific abstracts from PubMed that serves as the basis for the evidence-based validation.

In this context, our key objectives and contributions are:

1. **Comprehensive Framework:** To develop a comprehensive framework for automated fact-checking, specifically tailored to melatonin-related claims.
2. **Melatonin Specific Models:** We fine tuned existing state of the art language models for the melatonin corpus.
3. **User-Friendly Tool** To provide a user-friendly web-based tool for evaluating fact-checking algorithms, enabling end-users and researchers to assess the veracity of melatonin product claims using appropriate scientific evidence.

Beyond the immediate domain of melatonin-related claims, the ‘MelAnalyze’ framework has broader implications in being adapted to other domains as well. For instance, the ‘MelAnalyze’ framework can be easily extrapolated to evaluate other chemicals, mass consumer data, extracting insights into both positive and negative consumer experiences. Such insights can be useful in pharmaco-vigilance safety signal detection and in the triage of customer complaints. For consumer decision-making, recommendation systems significantly influence our choices^12^. However, there’s a risk of “algorithm over-dependence”, where consumers might place undue trust in these algorithm-generated recommendations, even when they might be flawed^13^. Coupled with aspects of human-computer interactions, persuasive technologies can influence purchasing decisions^14^. Together, their potential for disseminating misinformation or biased recommendations can have significant implications. By integrating a fact-checking layer, such as ‘MelAnalyze’, recommendation systems can provide more accurate and trustworthy suggestions. This ensures that consumers are not only receiving accurate product recommendations tailored to their preferences but also that these recommendations are grounded in verifiable facts. Ensuring that product claims and recommendations are validated against scientific evidence or verified facts can lead to enhanced consumer trust and better decision-making. We have illustrated some more of the impact areas in health in Figure 1. In summary, ‘MelAnalyze’ not only addresses the challenges of fact-checking melatonin-related claims but also showcases the broader applicability and necessity of automated fact-checking. In the subsequent sections, we delve deeper into background and related work, methods, results, and challenges.

**Figure 1.**
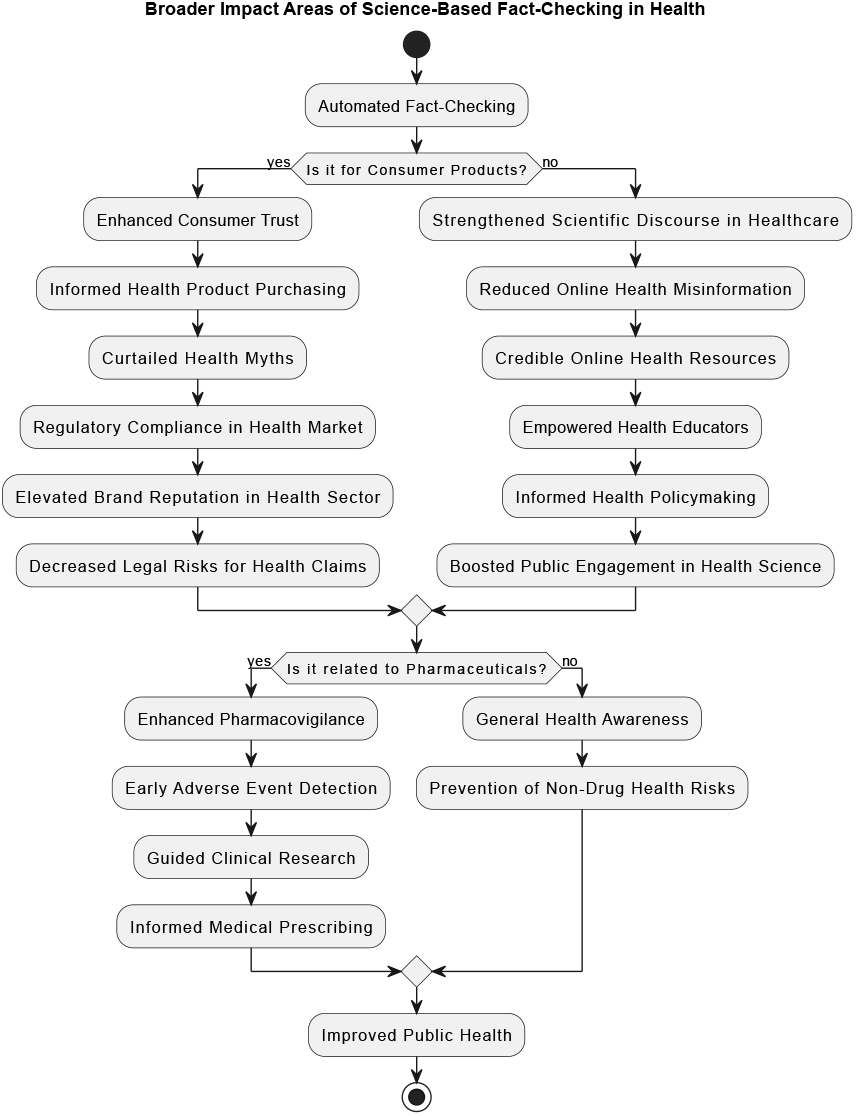
Overview of the broader impact areas of science based fact checking in health

**Figure 2.**
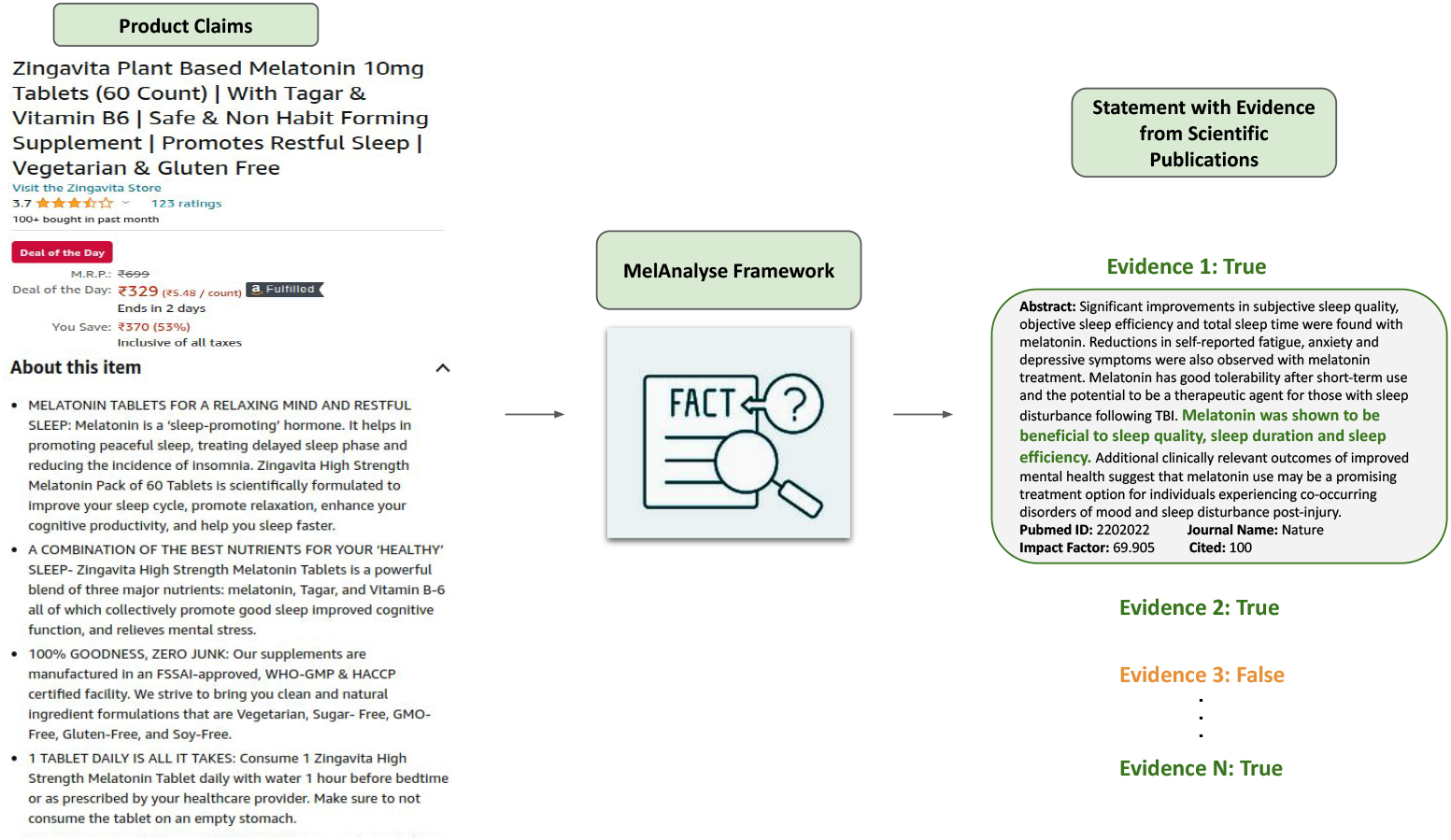
Problem Formulation

## 2 Background and related work

In this section, we will lay the foundational concepts that are used in the “MelAnalyze” framework. The intention is for it to be a primer of the different aspects, we refer the interested reader to get more detailed information from the corresponding references.

### 2.1 Natural Language inference

The Natural Language Inference (NLI) task consists of ascertaining the logical relationships between pairs of sentences. Given a premise sentence *P* and a hypothesis sentence *H*, the goal is to determine the nature of the relationship between them. Specifically, classifying if the relationship between *P* and *H* falls into one of three categories: “entailment,” “contradiction,” or “neutral”. “Entailment” signifies that the meaning of *H* can be logically inferred from *P*, indicating a stronger relationship between the two sentences. A “contradiction” indicates that *H* contradicts the information presented in *P*, reflecting a clear inconsistency. The “neutral” label implies that *H* neither entails nor contradicts *P*.

NLI models typically use a supervised learning approach. Given a labeled data set of premise-hypothesis pairs, the goal is to train a model that can generalize the relationship classification to unseen examples. The introduction of large-scale pretrained language models and its variants, has significantly advanced the state-of-the-art in NLI. The general approach involves pretraining the model on a massive corpus to learn contextualized representations of words and sentences. Subsequently, the model is fine-tuned on NLI-specific data sets to adapt its knowledge to the task at hand. During fine-tuning, the model learns to make accurate predictions. The model processes the premise and hypothesis sentences through its architecture and generates embeddings for both. These embeddings are then utilized for classification, often through a neural network layer. The final output layer provides the probabilities of the three classes (“entailment,” “contradiction,” and “neutral”), and the predicted label is chosen based on the highest probability.

### 2.2 Towards automatic fact checking

Fact checking involves the application of NLP techniques to automate the process of verifying the accuracy of claims, statements, or information. NLI serves as a fundamental framework for fact checking, enabling the assessment of the relationship between a claim and available evidence.

The utilization of NLI models for fact checking begins by inputting the claim into the system. The claim is transformed into their semantic representations, often in the form of embeddings as discussed before. Subsequently, the system compares this embedding against embeddings of potential evidence from reliable sources. Using cosine similarity as the similarity metric, the system ranks the evidence(s) based on their semantic proximity to the claim. Evidence(s) with smaller cosine distances are considered more relevant to the claim. The highest-ranked evidence(s) can then be presented to NLI algorithm, aiding them in making informed judgments about the claim’s veracity.

Automated fact checking have found applications in the domain of scientific evidence and bio-medicine. In the realm of scientific research and literature, NLI techniques have been employed to assess the compatibility of hypotheses with existing evidence, aiding researchers in formulating novel insights. It also plays a crucial role in combating misinformation in bio-medicine. It assists in verifying the accuracy of health-related claims, ensuring that medical advice and information disseminated to the public are evidence-based and reliable.

### 2.3 Transformers and large language models

In recent years, the introduction of transformers and large language models have boosted the field of NLP. These advancements have not only reshaped the landscape of NLP tasks but have also boosted Natural Language Inference (NLI) tasks with greater accuracy and efficiency.

Traditional approaches to NLP often relied on hand-crafted features and domain-specific knowledge, which limited their scalability and adaptability. The advent of transformers^15^ marked a paradigm shift in NLP. Transformers employ self-attention mechanisms to weigh the significance of different words in a sequence, enabling them to capture contextual relationships regardless of word order. This innovation unlocked the potential to model long-range dependencies and capture nuanced linguistic patterns. One of the well known breakthroughs in large language models is BERT (Bidirectional Encoder Representations from Transformers)^5^. BERT’s bidirectional training approach, in which the model learns from both left and right context, established new benchmarks across a range of NLP tasks. BERT’s success inspired subsequent models, including RoBERTa^16^, a robustly optimized BERT variant, and others like GPT-3^6^ which focused on generative tasks.

In the context of NLI, which involves determining the relationship between two sentences (premise and hypothesis), both generative and discriminative models can be explored. Generative models aim to generate the label ( entails, contradicts or neutral) when fine tuned, while discriminative models focus on predicting the label of the relationship between the hypothesis with respect to the premise. We formulate the fact checking problem as a NLI task. More details are shared in further sections.

### 2.4 Semantic similarity and vector search

Computing Semantic similarity is an important task in numerous NLP tasks, including NLI. In recent years, with advances in vectorization of text, cosine distance metrics are used to quantify the semantic similarity of text. Computing semantic similarity involves transforming textual data into a suitable vector representation ( or embeddings) that captures the underlying semantic meaning. Sentence-BERT^17^, which is based on BERT architecture is one of the good models for computing vector representations for sentences. In this process, each sentence *s* is encoded into a dense vector *v*_*s*_ using the BERT model. The vectors are then normalized to unit length to ensure that the magnitude of the vector doesn’t affect the similarity calculation. Given two sentences *s*_1_ and *s*_2_, their cosine similarity is computed using the dot product of their normalized vectors:

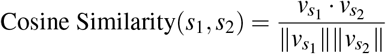

The model is trained with a contrastive loss function, encouraging similar sentences to be pulled together and dissimilar ones to be pushed apart in the vector space. This training strategy enables Sentence-BERT to capture intricate semantic nuances, thereby enabling more accurate semantic similarity comparisons. Once sentences are represented as vectors, the next step involves quantifying the semantic similarity between them. Cosine similarity is calculated as the cosine of the angle between two vectors and measures their similarity irrespective of their magnitude. The cosine similarity can be converted into a cosine distance by subtracting it from 1:

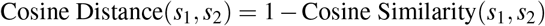

A smaller cosine distance implies a higher degree of semantic similarity between vectors, while a larger distance indicates greater dissimilarity. As such, cosine distance serves as an effective measure for ranking and identifying the most semantically similar sentences in NLI tasks. By generating semantically rich embeddings, Sentence-BERT enables the model to grasp subtle nuances in sentence meaning. This, combined with the cosine distance metric, empowers the system to quantify the semantic gap between sentences with precision. For vector search, the computed semantic similarity scores can be utilized to identify related sentences. The vector search efficiently retrieves relevant sentences from a database, which can then be used with NLI algorithms for the verification of textual claims.

### 2.5 Melatonin and its importance

Melatonin is primarily known for its role in regulating the sleep-wake cycle and was first discovered in the pineal gland^18–22^. However, the presence of melatonin has been detected in multiple extrapineal tissues and has garnered substantial attention in recent years due to its potential health benefits beyond sleep management^23^. Melatonin is widely used as a dietary supplement to address various sleep disorders, jet lag, and even certain neurological conditions. However, its popularity has also led to a surge in marketing claims, often making claims that go beyond the established scientific understanding^24^.

This is where fact checking comes in handy. With the help of a fact checking framework such as MelAnalyze, it becomes possible to scrutinize the information presented in marketing materials, product labels, and online content against reputable scientific sources. For instance, claims such as “Melatonin prevents all sleep disorders” or “Melatonin is completely risk-free” can be objectively assessed by fact checking algorithms against the existing body of biomedical literature. Moreover, the fact checking process can in principle also check details of melatonin’s mechanism of action, dosing recommendations, potential side effects, and interactions with other substances. This comprehensive evaluation ensures that consumers are provided with accurate and well-substantiated information, enabling them to make informed decisions about melatonin usage based on the best available evidence.

## 3 Data set preparation for Training and Validation

### 3.1 Data set for Model Training and Testing

The accuracy of our melatonin NLI models relies on the quality and diversity of the training data. To construct a robust and specialized NLI model for melatonin-related claims, a detailed data preparation process was undertaken. This section outlines the steps involved in curating the data sets. Details of the whole process is shown in the Figure 3.

**Figure 3.**
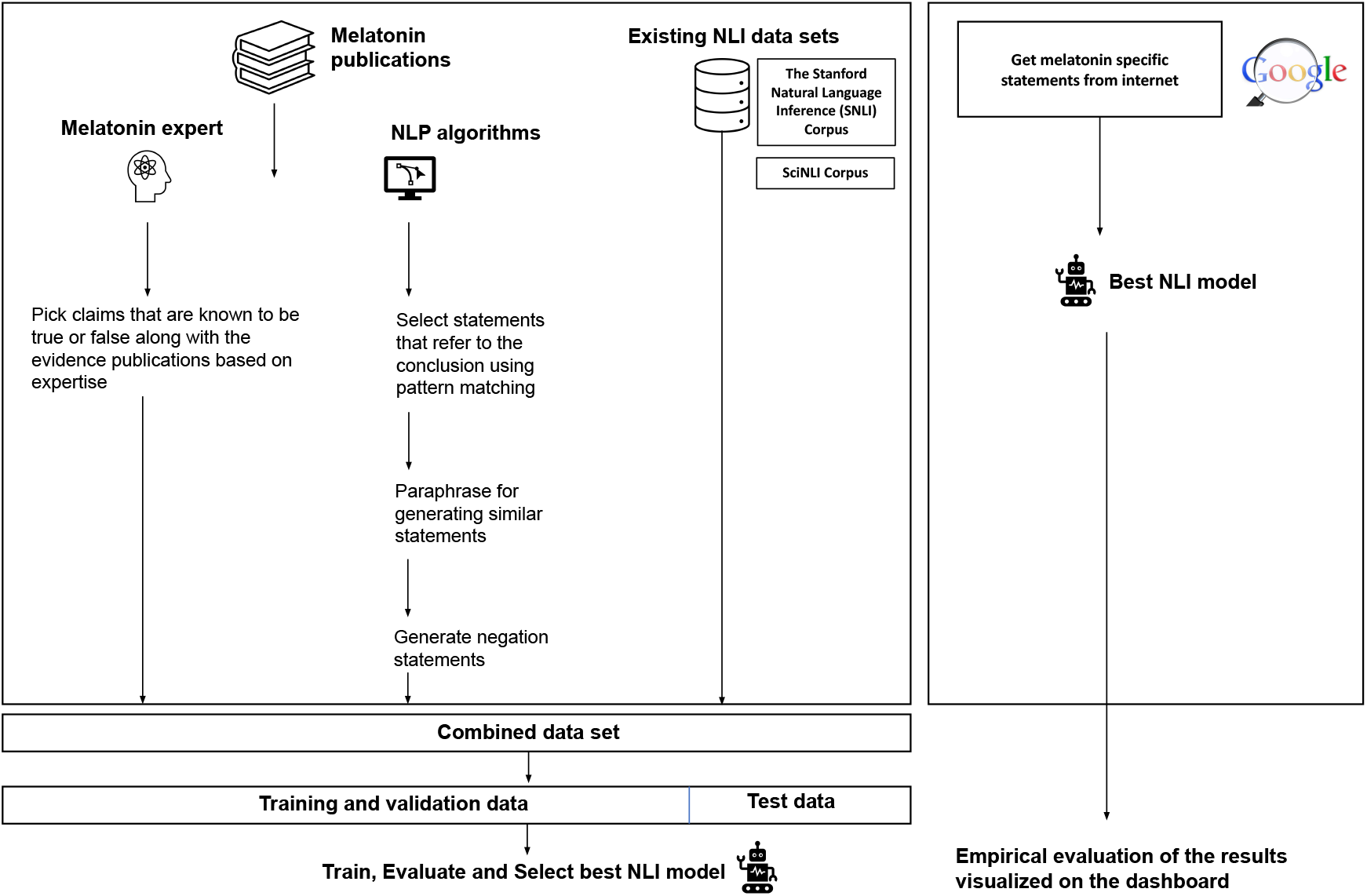
Overview of data-sets used for training the best NLI model for melatonin. The best NLI model was finally used on amazon product reviews of melatonin

#### 3.1.1 Expert-Guided Claim Selection

An important step involved collaborating with a melatonin expert to identify claims about melatonin that possess a definitive truthful status. These claims were selected based on the expert’s comprehensive knowledge of the field. Each claim was supported by established scientific evidence, thereby constituting a high-quality reference for training and validating the NLI model.

#### 3.1.2 Abstract-Based Statement Extraction

Building on the expert-verified claims, melatonin-related abstracts were extensively analyzed. Employing a pattern-matching strategy, statements corresponding to the abstracts’ conclusions were identified^25^. These statements served as the foundation for subsequent data augmentation and paraphrasing steps. To enhance diversity, a pre-trained NLP paraphrasing model was employed to generate paraphrased statement^26^ and negation statements^25^.

#### 3.1.3 Existing NLI data sets

To bolster the data set’s comprehensiveness, existing NLI data sets were incorporated. Specifically, the Stanford Natural Language Inference (SNLI) data set^27^ and the SciNLI data set^28^ were integrated. These data sets contributed a diverse range of general NLI instances, enriching the model’s ability to handle a wider spectrum of language structures and inferences. It is important to note that this data set was not melatonin specific.

#### 3.1.4 Combining data sets and Splitting for Training and Validation

The combination of these three distinct data sets resulted in a comprehensive training data set specifically tailored for melatonin-related NLI tasks. By combining the resulting training data encompassed specialized domain knowledge and also embraced the nuances of general NLI data sets. This combination aimed to strike a balance between domain-specific expertise and broader language understanding, fostering a versatile and well-informed melatonin-related NLI model. To facilitate effective model training and performance evaluation, a 80:20 split was employed.

### 3.2 Real-World claims data set for testing

The internet has become a significant platform for consumers to gather information about various products, including melatonin. Online forums serve as spaces where individuals share their experiences and insights regarding melatonin. We curated a data-set from the internet for validating the usefulness of our models. Data for this analysis was obtained by conducting a Google Search using the keyword “Melatonin medicine”. We collected information related to melatonin medicine from various sources, including blogs and articles. Next, we utilized the MelAnalyze Framework, which identified claims that were found false by the system and provided supporting evidence for their categorization. The Taxila^29^ tool was employed to streamline the collection procedure. This data set constituted real-world examples that could be subjected to empirical validation using the developed NLI model.

## 4 MelAnalyze framework

The “MelAnalyze” framework4 processes input claims through a series of steps. It begins with feature generation involving vector (embeddings) computation using the Sentence-BERT^17^ algorithm. Similarly, we compute embeddings from over 30,000 Melatonin-related PubMed abstracts which constitutes the knowledge base. Relevant sentences are fetched using semantic similarity, and the top 5 are shortlisted using both semantic and syntactic measures (cosine similarity). The best NLI model(s) is then applied to assess the input claim against these related sentences, generating multiple true/false or unclear assertions. Ultimately, the framework consolidates all these results and presents the finding via a user interface to the end user.

### 4.1 Natural language Inference models considered

The process of identifying the optimal NLI model for integration into the MelAnalyze framework included considering a combination of generative and discriminative models, as well as NLI-specific models. Generative models that were considered were LLaMA1 and LLaMA2, along with the discriminative model, RoBERTa. NLI-specific models, specifically `biobert_v1.1_PubMed_nli_sts`, `biobert-nli`, and `amoux/scibert_nli_squad`, were considered.

The process includes several key stages. First the NLI specific models are used as is in a zero shot setting to assess its effectiveness to be used out of the box. Next, each generative model is fine-tuned through prompt engineering, while RoBERTa is fine tuned using traditional fine tuning approaches. Subsequently, fine-tuning is performed on the NLI models too, yielding refined versions of the various models. Comprehensive evaluation is conducted to determine the efficacy of each fine-tuned model. This comprehensive assessment leads to the identification of the most suitable NLI model, which is then integrated into the MelAnalyze framework to facilitate accurate and effective claim evaluation. Figure5 discussed the evaluation of different NLI models. Details of the different models are shown below.

**Figure 4.**
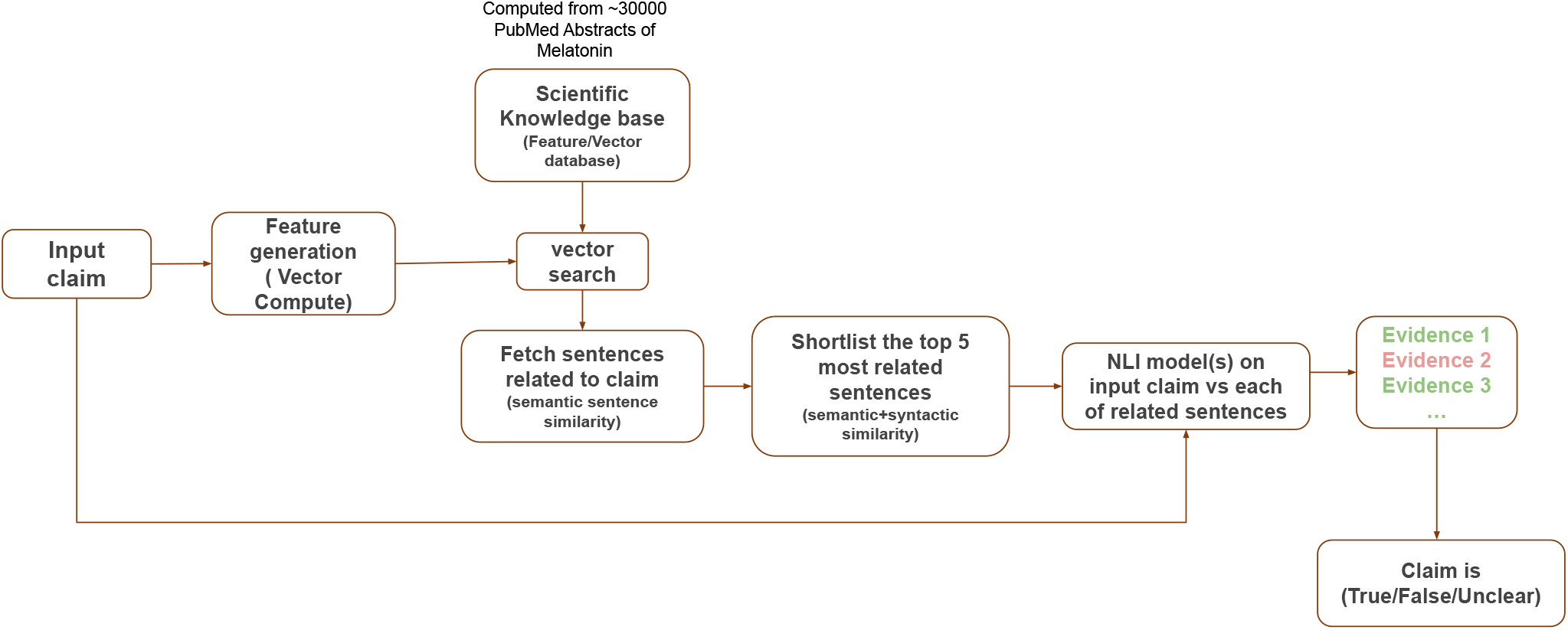
Overall framework of MelAnalyze

**Figure 5.**
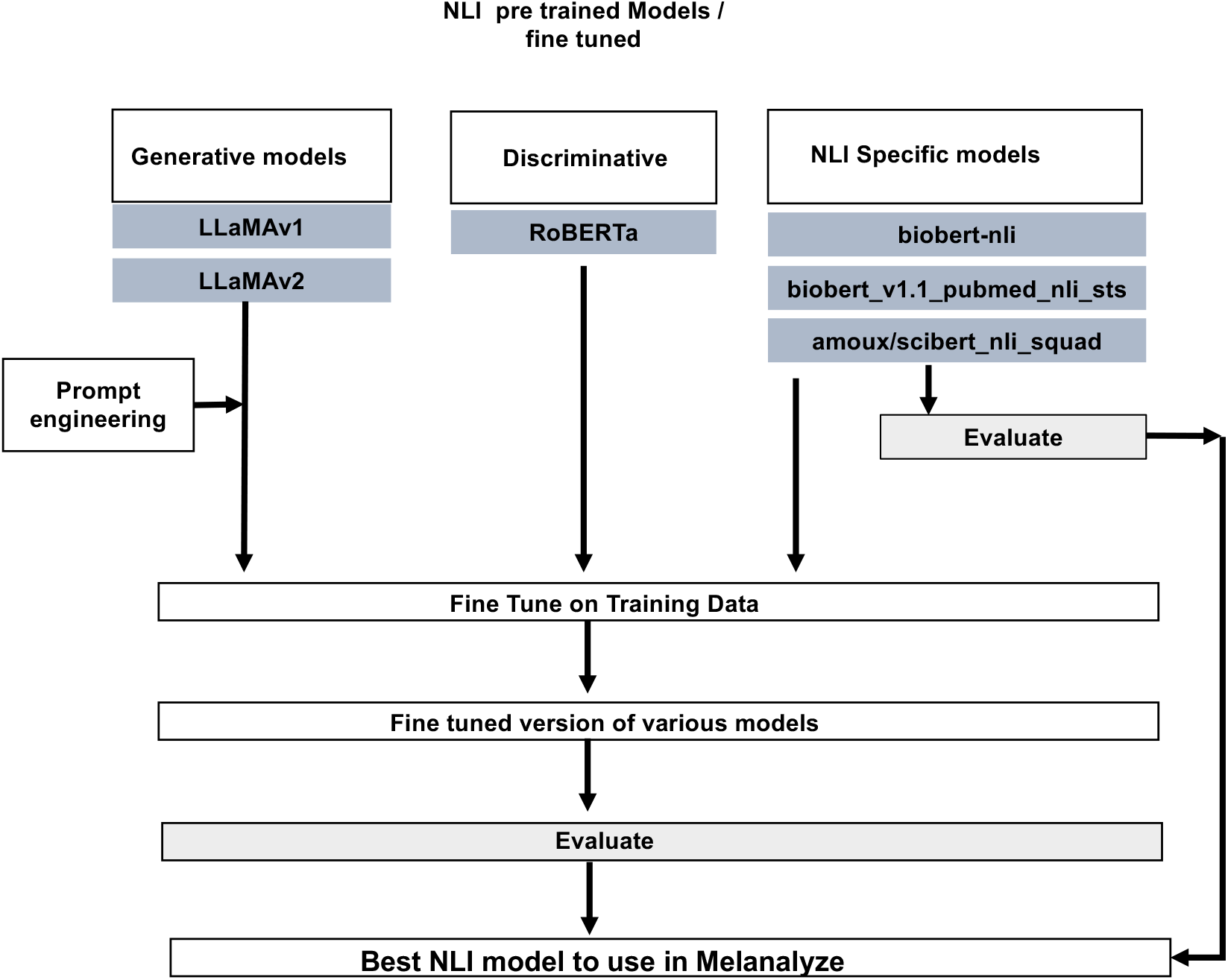
Overview of the different NLI Models considered and the fine tuning starategy applied

**Figure 6.**
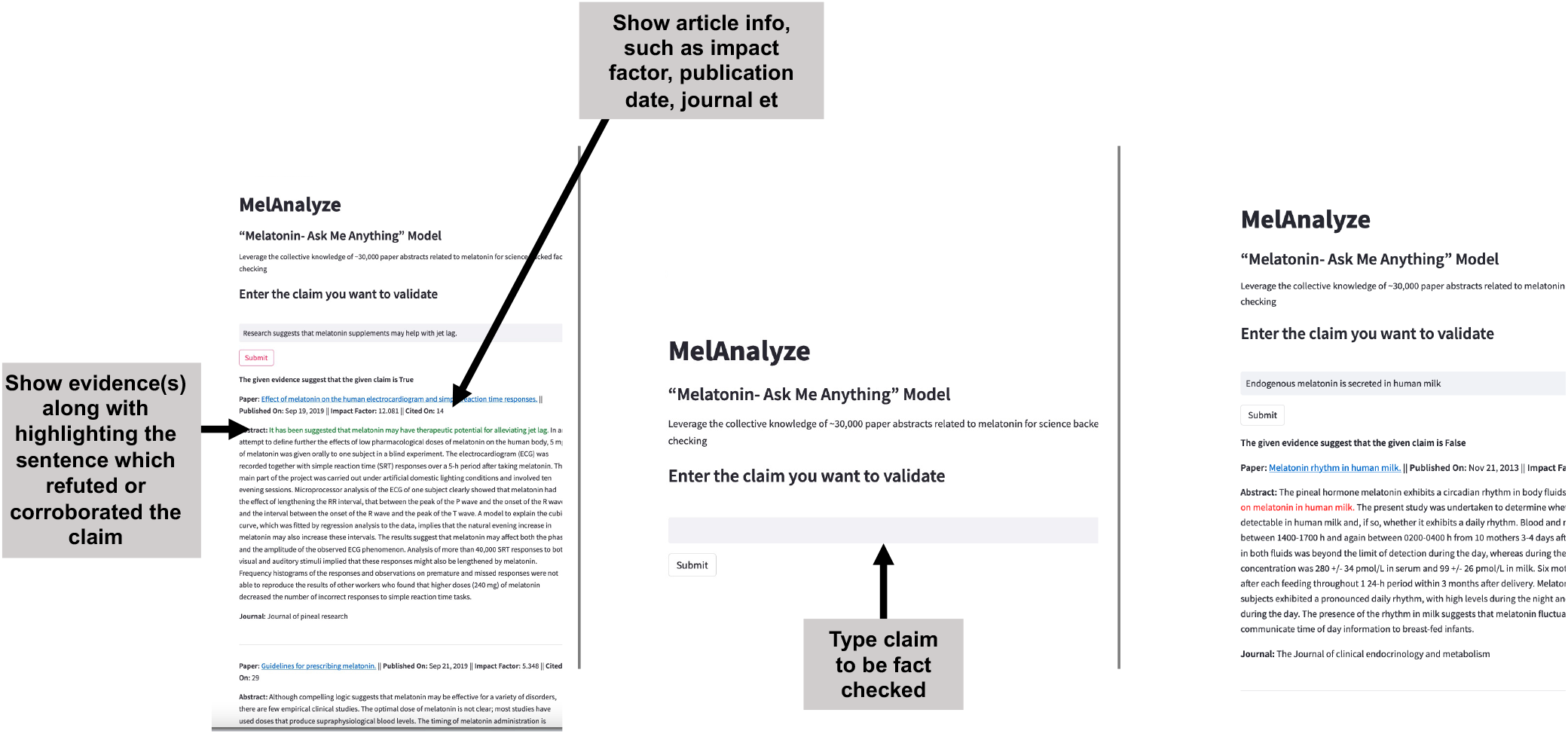
Overview of the Simple User interface for checking the veracity of claims using MelAnalyze. The center pane represents the main UI presented to the user. Once the user types the claim, in case the claim is true and backed by science (shown on left), the pertinent sentences in the abstracts are marked in green. Similarly if the claim is false ( shown in right), the sentences refuting the claim are marked in red

#### 4.1.1 LLaMA1

LLaMA1^30^ (Large Language Model Meta AI) is the firat generation of state-of-art foundational LLM designed by researchers at Meta and released on Feb 24, 2023. The LLaMA collection of language models ranges from 7 to 65 billion parameters in size, making it one of the most comprehensive language models. For this study, we have finetuned LLaMA1 under LORA^31^ setting which helps in low resource usage and is comparatively faster.

#### 4.1.2 LLaMA2

LLaMA 2^32^ is a collection of pretrained and fine-tuned large language models (LLMs) ranging in scale from 7 billion to 70 billion parameters. The architecture is very similar to the first LLaMA model, with the addition of Grouped Query Attention (GQA). The model is trained on 2 trillion tokens of data from publicly available sources. The pretraining setting and model architecture is adopted from LLaMA1. The training is done with QLORA^33^ (Quantization-aware Low-Rank Adapter Tuning) setup. QLoRA is a new technique to reduce the memory footprint of large language models during finetuning, without sacrificing performance.

#### 4.1.3 RoBERTa-base

RoBERTa^34^ (Robustly Optimized BERT Approach) is a variant of bert model, which was developed by Meta AI. It builds on BERT and modifies key hyperparameters, removing the next-sentence pre-training objective and training with much larger mini-batches and learning rates for longer duration. One key difference between RoBERTa and BERT is that RoBERTa was trained on a much larger data set and using a more effective training procedure. It achieved state-of-the-art performance on the MNLI, QNLI, RTE, STS-B, and RACE tasks.

#### 4.1.4 Biobert.v1.1.PubMed.nli.sts

Biobert based `biobert\_v1.1\_PubMed\_nli\_sts` model is a bert based binary model. This model is from Hugging-Face. It was trained on PubMed NLI data set. Since its trained on a PubMed scientific corpus, we chose this model for our experiments

#### 4.1.5 Biobert-nli

This is the model BioBERT fine-tuned on the SNLI and the MultiNLI data sets using the sentence-transformers library to produce universal sentence embeddings. The model uses the original BERT wordpiece vocabulary and was trained using the average pooling strategy and a softmax loss. `monologg/biobert\_v1.1\_PubMed` is the base model used for finetuning from HuggingFace’s AutoModel.

#### 4.1.6 Scibert.nli.squad

`Scibert\_nli\_squad` is another nli bert base model finetuned on scientific corpus. The model type is bert with 12 attention heads and 12 hidden layers. The max embedding size is 512. Since its trained on a scientific data set, we chose this model for our experiments

### 4.2 User interface: Web based tool

Our framework incorporates a user-friendly web-based interface as shown in Figure7 that serves as an intuitive platform for users to actively engage with the automated fact-checking process. This interface is designed to offer a seamless experience, allowing users to input their claims and receive real-time fact-checking results. A distinctive feature of our tool lies in its provision of comprehensive evidence derived from scientific publications. In addition to delivering the results, we furnish users with contextual information concerning the evidence sources. This includes key details like the impact factor of the journal where the evidence was published, comprehensive journal information, and the unique PubMed ID (PMID) linked to the respective publication. This enriched presentation of evidence not only facilitates the evaluation of claim but also empowers users to assess the credibility and reliability of the underlying sources. By presenting this supplementary information, our interface offers users a holistic view, allowing them to make well-informed decisions based on both the fact-checking outcomes and the supporting scientific evidence. This thoughtful design promotes transparency, encourages thorough examination, and contributes to a more responsible approach to navigating and disseminating information.

**Figure 7.**
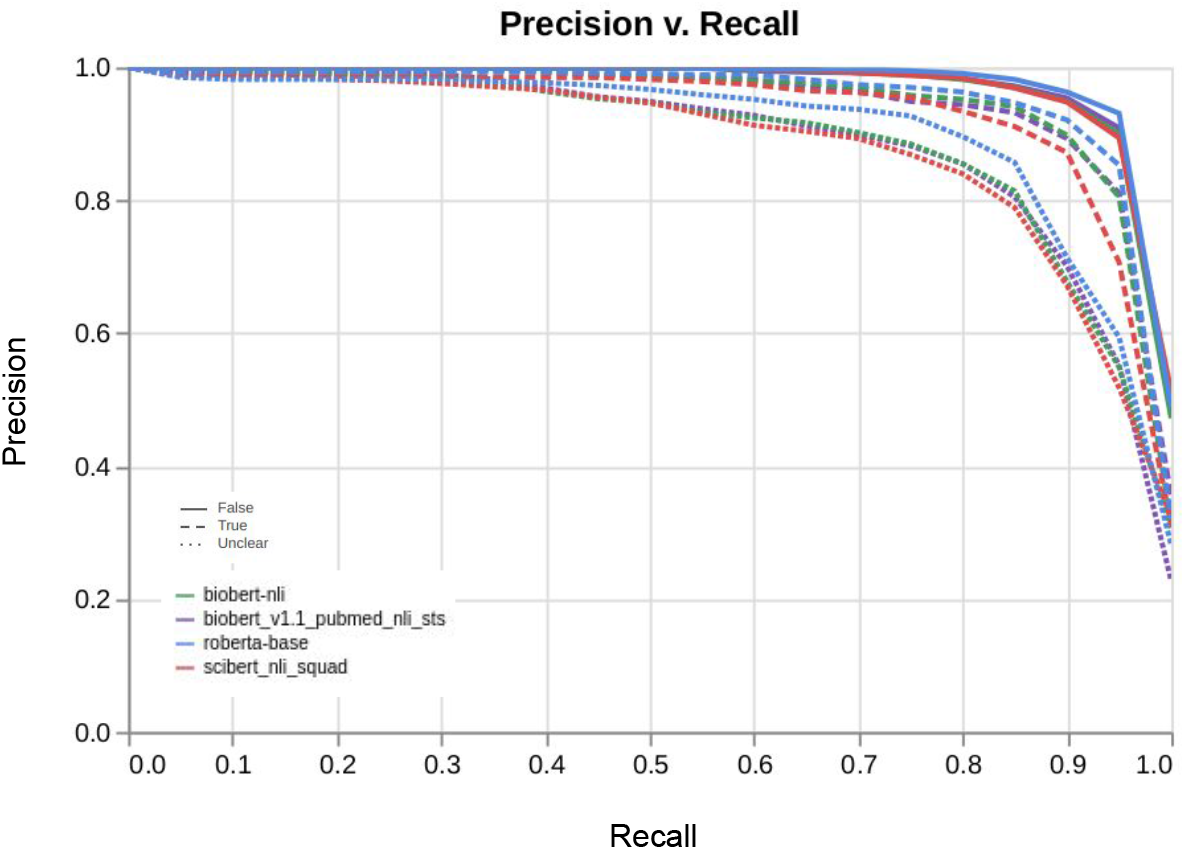
Precision recall curve for the various fine-tuned pre-trained models

## 5 Experimental results

The Table 1 presents a comprehensive overview of the experimental results obtained from different fine-tuned NLI models made for melatonin within the MelAnalyze framework. These models are classified into two main categories: generative models and discriminative NLI-specific models. The performance of each model is assessed using three fundamental metrics: precision, recall, and F1-score, which collectively offer a comprehensive evaluation of their effectiveness.

**Table 1.**
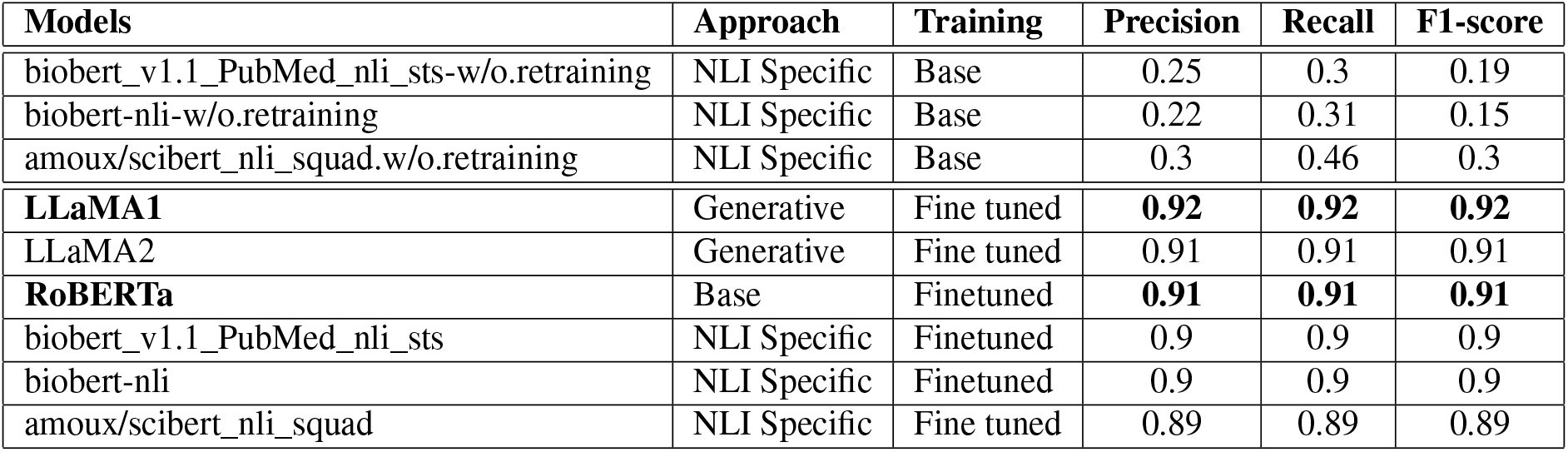
Performance Metrics of Various Fine Tuned and NLI Models. Models fine tuned on LLaMA1 and RoBERTa perfomed the best.

In the generative models category, LLaMAv1 exhibited remarkable performance, achieving a precision of 0.92, recall of 0.92, and an F1-score of 0.92. LLaMAv2 also demonstrated competitive results, with a precision of 0.91, recall of 0.91, and an F1-score of 0.91. In the discriminative NLI-specific models category, the RoBERTa model displayed consistent precision, recall, and F1-score values, all set at an impressive 0.91. Similarly, both the `biobert_v1.1_PubMed_nli_sts model` and the `biobert-nli` model showcased robust performance, maintaining precision, recall, and F1-score values of 0.9 across the board.

Furthermore, the table highlights the performance of models in their base and fine-tuned iterations. Notably, the base version of `biobert_v1.1_PubMed_nli_sts` yielded lower precision, recall, and F1-score values of 0.25, 0.3, and 0.19, respectively. Similarly, the base version of `biobert-nli` achieved relatively lower precision, recall, and F1-score values of 0.22, 0.31, and 0.15, respectively. The base version of `amoux/scibert_nli_squad` too showed poor results. In contrast, all fine-tuned NLI models demonstrated competitive metrics with increased accuracy highlighting the power of fine-tuning the NLI models with new melatonin-specific data. Among these models, the ones with the highest performance are highlighted in bold (corresponding to LLaMAv1 and RoBERTa fine-tuned), underscoring its good capability to effectively assess the veracity of claims through the MelAnalyze framework.

### 5.1 Experimental Parameters

We fine-tuned two generative models, namely LLaMA1-7B and LLaMA2-13B. For the LLaMA1 modes, fine-tuning was performed using the Instruction-tuning approach with LoRA setup. The tuning parameters details are as follows: `learning_rate` = 3 *×* 10^*−*4^, `batch_size` = 128, `micro_batch_size` = 4, `warm_iters` = 100. The LoRA setup was kept default with `lora_r` = 8, `lora_alpha` = 16, and `lora_dropout` = 0.05. The maximum sequence length was set to 512. For LLaMA instruction tuning, the Lit-LLaMA repository was employed. The LLaMA2 model was fine-tuned using the QLORA approach, which aims to reduce the memory footprint of large language models during fine-tuning without compromising performance. The LoRA configuration based on the QLORA paper is as follows: `lora_alpha` = 16, `lora_dropout` = 0.1, `r` = 64, `bias` = “none”, `task_type` = `“CAUSAL_LM”`. Other parameters include `num_epochs` = 10, `batch_size` = 32, `gradient_accumulation_steps` = 2, `optim` = `“paged_adamw_32bit”`, `learning_rate` = 2 *×* 10^*−*4^, `bf16` = True, `tf32` = True, `max_grad_norm` = 0.3, and `warmup_ratio` = 0.03. Both models were trained on an NVIDIA A40 GPU instance. LLaMA took 1.5 days to train, while LLaMA2-13B was completed in 5days. Both models utilized around 31GB of GPU memory during training.

For the fine-tuning of BERT-based model Roberta and the other three SCI-NLI models, the following parameters were used: `num_epochs` = 5, `train_batch_size` = 8, and `learning_rate` = 1 *×* 10^*−*5^. All BERT models were trained on an NVIDIA GeForce RTX 2080 Ti with 12GB of memory. The training process for each model took approximately 5-6 hours, utilizing around 5GB of GPU memory.

### 5.2 Empirical evaluation of the MelAnalyze framework on claims on the internet

Here are several claims gathered from the internet and assessed by the MelAnalyze system. The claims listed below have been categorized as “False” by the system. Additionally, there are claims categorized as “True” included in the following list.

**Claim:** Doctors might prescribe melatonin for children with additional needs, but the Therapeutic Goods Administration has not approved melatonin for use by typically developing children.

**Evidence:** The data support the notion that melatonin, or one of its analogs, might find use as an anesthetic agent in children.

**Verdict:** False

**Claim:** The American Academy of Sleep Medicine recommends against using melatonin because the supplement’s side effects are a serious concern in this group of people.

**Evidence:** The American Academy of Sleep Medicine recommends the timed use of the chronobiotic melatonin to hasten adaptation.

**Verdict:** False

**Claim:** Melatonin should not be used as a substitute for good sleep hygiene and consistent bedtime routines in children.

**Evidence:** In children as well, melatonin has value as a sleep-promoting agent.

**Verdict:** False

**Claim:** Melatonin aggravates ulcerative colitis

**Evidence:** Melatonin ameliorated ulcerative colitis-associated local and systemic damage in mice.

**Verdict:** False

**Claim:** Melatonin increases the destruction of articular cartilage and bone in rheumatoid arthritis.

**Evidence:** Melatonin (MLT) can increase the expression of cartilage-derived growth factor and stimulate the synthesis of cartilage matrix.

**Verdict:** False

**Claim:** Specific inhibitors of melatonin would benefit patients with asthma.

**Evidence:** We conclude that melatonin can improve sleep in patients with asthma.

**Verdict:** False

**Claim:** Melatonin is associated with enhanced oxidative stress and inflammation status in multiple sclerosis

**Evidence:** In addition, melatonin is associated with decreased oxidative stress and increased anti-oxidative factors.

**Verdict:** False

**Claim:** Being exposed to light at night can block melatonin production.

**Evidence:** Ocular light exposure at night can suppress melatonin levels in humans.

**Verdict:** True

**Claim:** Research suggests that melatonin supplements may help with jet lag.

**Evidence:** It has been suggested that melatonin may have therapeutic potential for alleviating jet lag.

**Verdict:** True

**Claim:** The American College of Physicians guidelines strongly recommend the use of cognitive behavioral therapy for insomnia (CBT-I) as an initial treatment for insomnia.

**Evidence:** Cognitive behavioural therapy for insomnia is recommended as the first-line treatment for chronic insomnia in adults of any age (strong recommendation, high-quality evidence)

**Verdict:** True

**Claim:** Melatonin, a full service anti-cancer agent: inhibition of initiation, progression and metastasis.

**Evidence:** Melatonin is a potent oncostatic agent and it prevents both the initiation and promotion of cancer.

**Verdict:** True

**Claim:** Melatonin reduces night blood pressure in patients with nocturnal hypertension.

**Evidence:** In patients with essential hypertension, repeated bedtime melatonin intake significantly reduced nocturnal blood pressure

**Verdict:** True

## 6 Conclusion and future directions

In summary, the combination of NLI models, semantic similarity, and automated fact-checking has enabled an effective approach for evaluating scientific claims, specifically for melatonin-related information. Utilizing advanced NLI models, we have demonstrated the feasibility of this method in assessing claim accuracy and combating misinformation. Looking forward, integrating more refined NLI-specific models and tailored training data can enhance the precision of fact-checking. Additionally, incorporating contextual features and linguistic nuances could further improve the MelAnalyze capabilities for nuanced claim analysis. Beyond melatonin, our approach has broad applications across diverse domains.

## 7 Data and Tool Availability

All data and tool information is available at the supplementary companion website https://bit.ly/melanalyze_tool

## 8 Acknowledgements

The authors acknowledge the funding provided by Vectura Fertin Pharma Inc., for a portion of this study. The authors thank Dr. Julia Hoeng and Gordon Dawson from Vectura Fertin for their strategic advice and Guidance through out the course of this study. This work was originally done in October 2023.

## 9 Author contributions statement

SKP, SG, and GE Conceived the study. SKP designed the solution strategy. GE prepared the expert curated dataset. NK conducted the experiments and generated the results. All authors wrote and reviewed the paper.

